# Urban birdsongs: higher minimum song frequency of an urban colonist persists in a common garden experiment

**DOI:** 10.1101/761734

**Authors:** Dustin G. Reichard, Jonathan W. Atwell, Meelyn M. Pandit, Gonçalo C. Cardoso, Trevor D. Price, Ellen D. Ketterson

## Abstract

Environmental changes caused by urbanization and noise pollution can have profound effects on acoustic communication. Many organisms use higher sound frequencies in urban environments with low-frequency noise, but the developmental and evolutionary mechanisms underlying these shifts are less clear. We used a common garden experiment to ask whether changes in minimum song frequency observed 30 years after a songbird colonized an urban environment are a consequence of behavioral flexibility or canalized changes that occur early in development. We captured male juvenile dark-eyed juncos (*Junco hyemalis thurberi*) from two recently diverged populations (urban and mountain) soon after they reached independence (aged 25-40 days), raised them in identical indoor aviaries, and studied their songs at an age of three years. We found that the large population difference in minimum frequency observed in the field persisted undiminished in the common garden despite the absence of noise. We also found some song sharing between the common garden and natal field populations, indicating that early song memorization before capture could contribute to the persistent song differences in adulthood. These results are the first to show that frequency shifts in urban birdsong are maintained in the absence of noise by genetic evolution and/or early life experiences.

## Introduction

Anthropogenic noise can alter the biology of diverse animal taxa at organismal, population, and even community scales [1–7]. In particular, the low frequency background noise often associated with urbanization can interfere with animal communication and has been associated with changes in acoustic signals that improve sound transmission [8, 9]. One such change that is widely observed in urban environments is increased minimum frequency of acoustic signals, which may be an adaptation to overcome the masking effects of low-frequency noise [10–16]. Our understanding of the developmental and evolutionary mechanisms that may underlie such changes in acoustic signaling remains limited [9, 17–22]. Species such as oscine songbirds that learn their songs are of particular interest due to the potential for cultural evolution and other forms of behavioral plasticity, which can facilitate rapid change in response to anthropogenic noise [23–25].

Several non-mutually exclusive hypotheses have been proposed to explain changes in song frequency in urban environments [9, 26], including short-term plasticity, ontogenetic effects (early experience), and evolutionary change across generations.

The plasticity hypothesis argues that frequency shifts are the result of behavioral flexibility in response to the presence or absence of a noise stimulus. Some studies in oscine songbirds have found evidence supporting plasticity either through rapid increases in minimum song frequency [18, 22, 27, 28] or switching to song types with higher minimum frequencies when noise is present [17]. However, evidence from other songbird species has indicated that short-term plasticity in response to noise does not always explain the frequency shifts present in urban birdsong [20, 29–32].

The early experience hypothesis argues that the presence of noise during development affects song structure and production later in life. For example, evidence from black-capped chickadees (*Poecile atricapillus*) and white-crowned sparrows (*Zonotrichia leucophrys*) suggests that experiencing noise early in life may be necessary for the development of noise-induced plasticity in song frequency in adulthood [33, 34]. In addition to inducing plasticity, noise during development may mask lower frequency tutor songs and cause selective learning of songs with higher minimum frequencies in urban environments [35]. Evidence suggests that some species preferentially learn songs that are less degraded by environmental transmission [36, 37; but see 19]. Notably, a recent study in white-crowned sparrows found that males developing in an environment with low-frequency noise preferentially learned higher frequency (less masked) songs [21]. Collectively, these results indicate that ontogenetic effects of experiencing noise during early life may affect song frequency in adulthood by the preferential learning of certain songs (i.e. cultural selection), or by developing the ability for plastic adjustments to noise.

Finally, the evolutionary change hypothesis argues that natural or sexual selection or drift on relevant genetic variation influences song frequency across generations [9, 26]. In this case, the likely selective pressure is the masking of low-frequency songs by anthropogenic noise, making those songs less adaptive (e.g. less effective signals for territoriality and mate attraction) [21]. As a result, individuals with cognitive, morphological, or sensory phenotypes that cause them to learn and/or produce higher frequency songs will have higher fitness leading to directional selection [38]. The colonization of urban environments is associated with a diverse array of phenotypic divergence, such as changes in body size [39, 40], bill morphology [41], and neural architecture [42], some of which is likely to be genetic, and all of which may contribute to evolution in song structure.

Here, we used a common garden experiment to test predictions of these hypotheses. Common garden experiments are a powerful method for differentiating between the relative effects of genetic and environmental factors in determining phenotypic differences. The only common garden study of divergence in urban acoustic signals thus far used a species of grasshopper (*Chorthippus biguttulus*) [14]. In that study, individuals originating from noisy environments sang at significantly higher minimum frequencies than individuals from quiet environments, but individuals reared in a noisy common garden environment produced higher frequency songs as adults regardless of their population of origin. Collectively, these results suggest roles for both evolutionary change and early noise exposure in determining the differences in song frequency of urban populations. It is not known whether a similar interplay of evolutionary and ontogenetic effects applies to birdsong, which can be learned culturally, and should show much greater plasticity in response to noise than the stridulatory songs of insects [43].

We studied an urban population of dark-eyed juncos (*Junco hyemalis thurberi*) that was recently established in the early 1980s [44, 45]. This population ceased migrating and rapidly diverged in a variety of behavioral, hormonal, and life-history traits from an ancestral, migratory population that breeds in a wildland environment in the inland mountains and migrates to the coast during the winter [19, 46, 47]. The urban and mountain acoustic environments differ strongly, including in anthropogenic noise, which is negligible in the mountains [48], and urban juncos sing at significantly higher minimum frequencies (ca. 0.5 kHz higher) than juncos in the mountain population [19, 48, 49]. If the higher minimum frequency observed in the urban population is caused by behavioral flexibility in song production (plasticity), then the population difference should disappear in the common garden. Alternatively, if the population difference is genetically-based or is affected by early experience, including song learning, then the population difference should persist in the common garden. To assess whether early song learning can account for population differences in the common garden, we compared song types of the common garden birds to songs from their natal populations, searching for cases of song sharing.

## Methods

The details of our study populations and the common garden experiment are reported elsewhere [46, 47]. Briefly, we studied two populations of dark-eyed juncos (*Junco hyemalis thurberi*) in San Diego County, California, USA. The urban population was located at the University of California, San Diego (hereafter “urban”; elevation 30 m; lat. 32°52’N, long. 117°10’W) and the wildland population was located at Laguna Mountain Recreational Area (hereafter “mountain”; elevation 1,700 m; lat. 32°52’N, long. 116°25’W). In July 2007, we captured 80 independent, juvenile juncos (day 25-40 post-hatch as determined by prior banding in the nest for most subjects, see [47]) from the urban and mountain populations (20 per sex per population). The birds were transported to Indiana University in Bloomington, Indiana, USA, and housed in mixed sex flocks in separate but identical indoor aviaries (6.4 x 3.2 x 2.4 m) with light conditions that mimicked seasonal shifts in their native ranges [47]. The two populations were acoustically isolated from one another such that males could only hear and interact with members of their own population. The current study took place in May 2010, which was the third spring that these males experienced in the common garden.

### Song Recordings

Dark-eyed juncos produce a loud, broadcast song consisting of a simple trill (rapid repetition of the same element), and each male sings a small repertoire of distinct song types (Fig. 1; [50]). To record songs, we isolated each surviving male from the common garden (mountain, *N* = 10; urban, *N* = 8) in a (45.7 x 45.7 x 45.7 cm) cage with a single perch extended across its center and access to food and water *ad libitum*. An Audio-Technica shotgun microphone (Model AT835b) was suspended ~30 cm above the center of the cage’s perch and connected to a Marantz digital recorder (Model PMD660). We recorded each male for three hours using a 44.1 kHz sampling rate in uncompressed WAV format. One mountain male did not sing and was excluded from the analysis.

**Figure 1.**
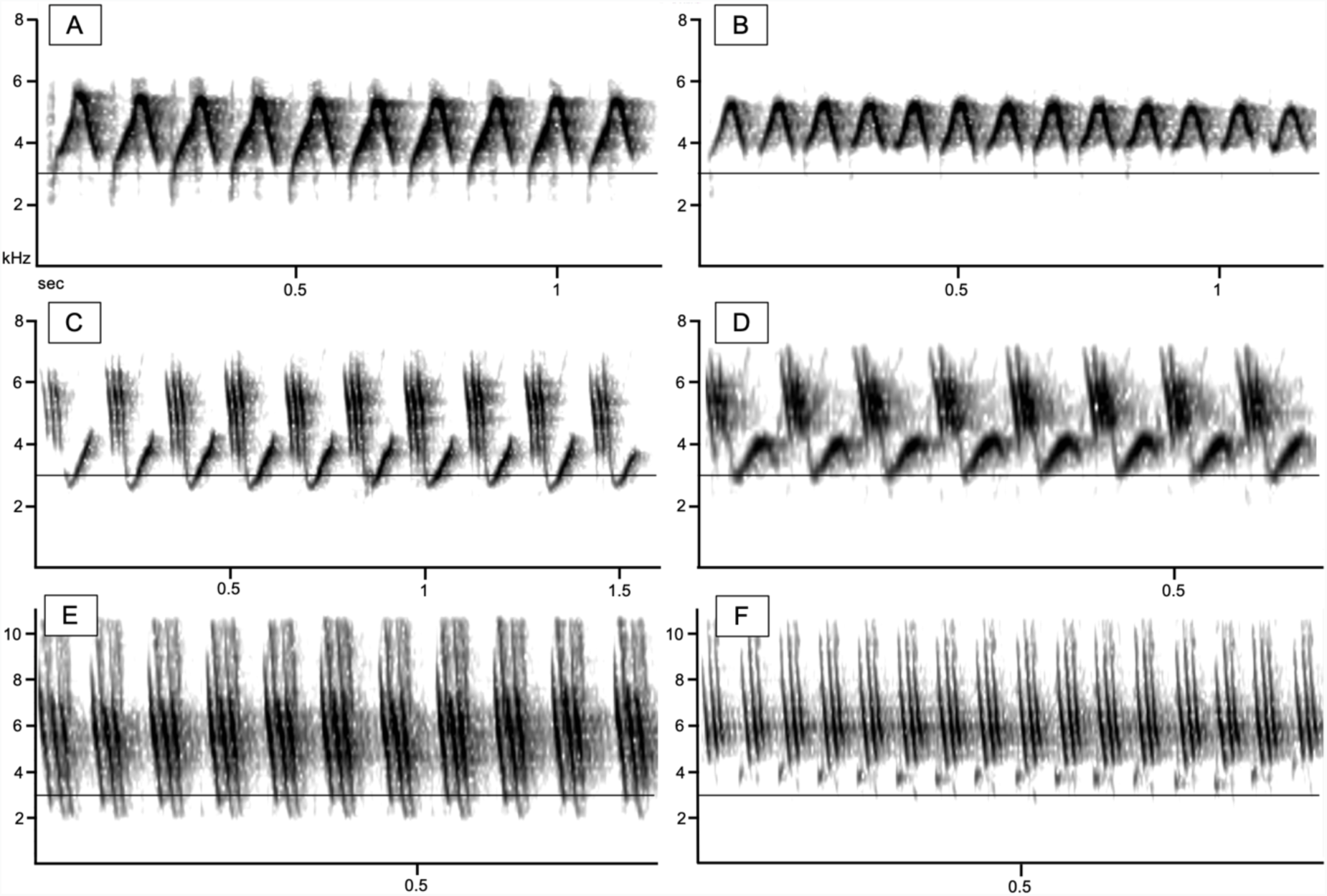
Example spectrograms of shared song types observed between mountain- and urban-captured males in the common garden. Songs A, C, and E were produced by mountain males and songs B, D, and F were produced by urban males. The thin line on each spectrogram marks 3 kHz to highlight frequency shifts between mountain and urban songs.

Junco song types can be reliably distinguished by visual comparisons of spectrograms [49, 51, 52]. Assignments of song types and song type sharing were performed blindly without any knowledge of the population of origin and were confirmed by at least two independent observers. We recorded 17 different song types from the mountain population, some of which were shared by more than one bird (see Results), and 28 different song types from the urban population, some also shared. Mountain males sang an average of 3 song types (range: 1-5), and urban males sang an average of 4.9 song types (range: 2-7). We also compared these songs to a catalog of song types from both field populations collected in 2006 and 2007 (mountain, *N* = 115; urban, *N* = 168; [19]) to assess sharing between the common garden males and their natal populations.

### Song Measurements

We used Raven Pro 1.4 [53] to measure minimum, maximum, and peak frequency. Measurements were performed by the same observer (MPP) who was blind to population of origin to avoid bias. For each combination of song type and male, we randomly selected a representative exemplar and generated a spectrogram (Hann Window, 512 DFT, 86.1 Hz frequency resolution, 5.8 ms time resolution). We used the cursor to visually draw a “selection box” around each song type, bounded by the perceived start and end time as well as the minimum and maximum frequency. We recorded the minimum and maximum frequencies of these visual measurements from the spectrogram, and also recorded the peak frequency of the selection (frequency with the highest cumulative amplitude).

Visual measurements from spectrograms have been criticized as a potentially biased technique for determining minimum and maximum frequency [54–57]. Instead, researchers have advocated using the power spectrum and a threshold value as a more objective alternative. To assess potential differences in these techniques, we also measured minimum and maximum frequency from the power spectrum of each song using a threshold of minus 30 dB relative to the peak frequency of the song. Minimum and maximum frequency were defined as the points of intersection between the power-spectrum curve and the threshold value [55, 58]. We were able to use a very large amplitude threshold (minus 30 dB) because of the high signal-to-noise ratio in our aviary recordings. Nonetheless, in a few song types, faint harmonics caused power-spectrum measurements of maximum frequency far exceeding the highest fundamental frequency observed on the spectrogram and the normal range of maximum frequencies measured in various field studies [50]. Similarly, there was one song type where the power-spectrum measurement of minimum frequency resulted in a value that was much higher than the lowest observed frequency. We excluded those cases (*N* = 1, minimum frequency; *N* = 10, maximum frequency) from any analyses involving the power-spectrum measurements for maximum frequency.

All raw frequency measurements were log_10_ transformed before further analysis, because perception and modulation of sound frequency both function on a ratio scale [59]. Log transformation facilitates the comparison of frequency differences across different frequency ranges; otherwise, differences in maximum or peak frequency would be over-estimated compared to differences in minimum frequency.

Across all measurements of minimum frequency, the visual measurements from spectrograms and threshold measurements from power spectra were significantly correlated (*r* = 0.79, *N* = 64, *P* < 0.001; Fig. S1A), and there was only a slight, but statistically significant difference in their means (0.028 log_10_Hz [168.7 Hz]; *t*_63_ = −3.49, *P* < 0.001; Figure S1). Maximum frequency measurements were also correlated across methods (*r* = 0.96, *N* = 55, *P* < 0.001; Fig. S1B), and their means did not differ significantly (*t*_54_ = −1.85, *P* = 0.07; Fig. S2). In the main text, we only report analyses using visual frequency measurements from spectrograms to facilitate a comparison with a dataset of field recordings previously analyzed in this manner [see above; 19]. In the supplementary material we report alternative comparisons between the common garden populations using threshold measurements from power spectra (Tables S1 and S3), which yielded identical results to those reported in the main text.

### Statistical Analysis

To compare acoustic traits between populations in the common garden, we conducted linear mixed models (LMM) using the lme4 package in R version 3.5.2 [60, 61]. Each model contained a song measurement as the response variable, population of origin as an independent factor, and song type as a random factor. Song types were used as a random factor in the main text, rather than male identity, because junco song traits are a property of the individual song type (high within-type repeatability across males) rather than a property of the individual males [low repeatability across song types in the repertoire of individual males; 49]. In the supplementary material we report identical analyses using male identity as a random effect, instead of song type, and our results remain unchanged (Tables S2, S3).

To assess whether early song learning in the natal urban environment, as opposed to songs that developed later in the common garden, was important to maintain high minimum song frequency in urban-captured males, we compared the minimum frequency of song types shared with urban field recordings and song types not shared with field recordings. Since junco song development is strongly influenced by social learning and by creating novel song types (see below; reviewed in [50]), comparing shared and non-shared song types can test whether social learning influences acoustic traits in a particular direction [62]. We used a LMM with minimum frequency as the dependent variable, song type as a random factor and whether song types were shared or not with urban field recordings as an independent factor. Finally, we used *t*-tests to compare frequency measurements from the common garden with those from a previously published field study of both populations [see above; 19].

## Results

We identified 7 song types (out of 17 total song types; 41%) that were shared between two or more mountain-captured males in the common garden, and 8 song types (out of 28; 29%) that were shared between two or more urban-captured males. Three song types were shared between populations in the common garden (Fig. 1). We also identified 2 song types from the mountain-captured males (out of 17; 12%) that had been previously recorded from mountain males in the field (out of 115 song types recorded in the field), and 6 song types from the urban-captured males (out of 28; 21%) that had been previously recorded from urban males in the field (out of 168 song types recorded in the field).

In the common garden, males that were captured in the urban population sang with significantly higher minimum frequencies than mountain-captured males (*t* = 3.59, *P* < 0.001; Fig. 2 and examples in Fig. 1). In contrast, we found no detectable differences between the common garden populations in maximum (*t* = −0.38, *P* = 0.71) or peak frequency (spectrogram: *t* = 0.31, *P* = 0.75).

**Figure 2.**
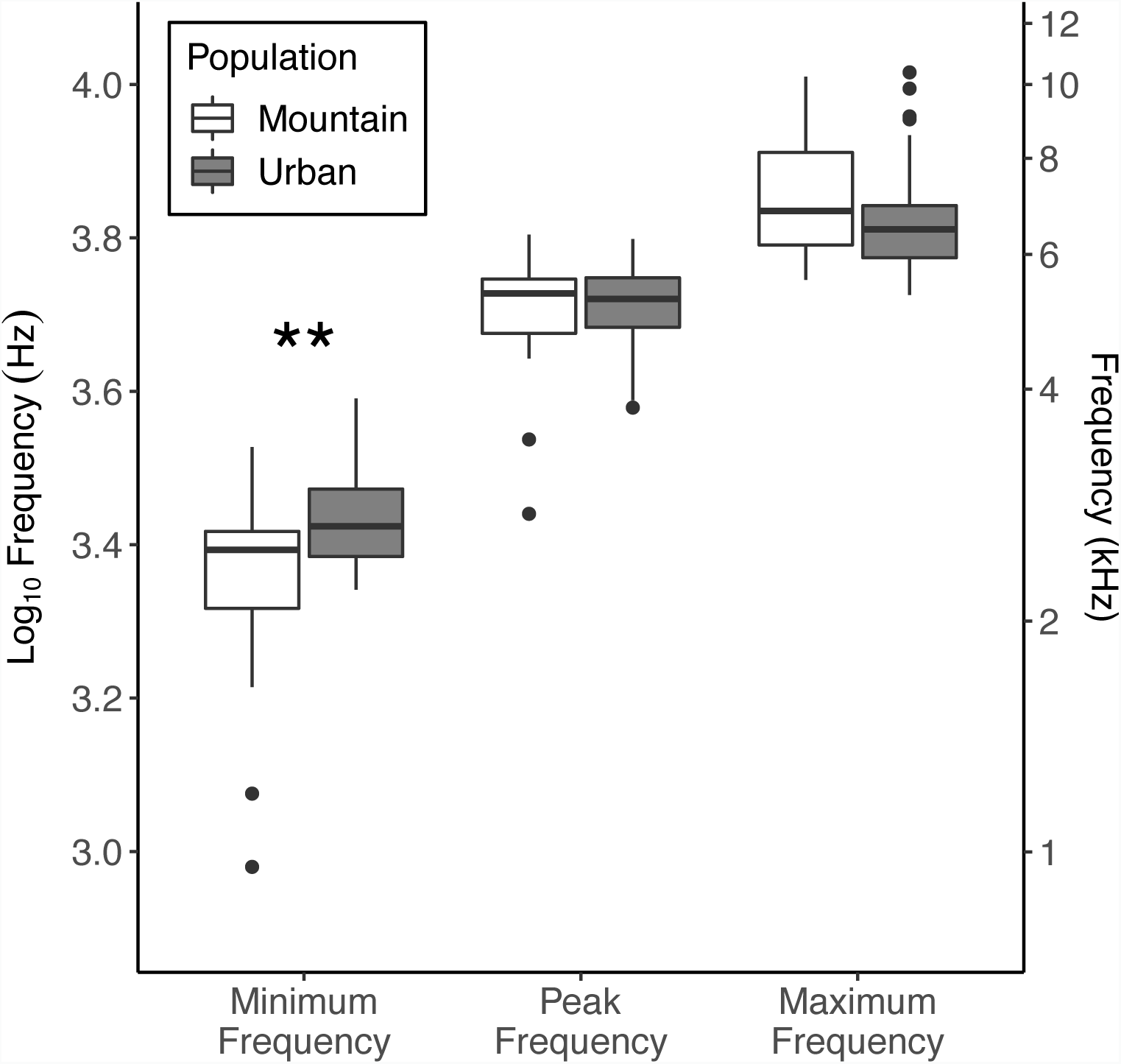
Comparison of the minimum, peak, and maximum frequencies of song types produced by mountain- and urban-captured males raised in a common garden environment. Each box represents the interquartile range and median, whiskers represent range of data within 1.5 times the interquartile range, and dots represent data points exceeding that range. \*\**P* < 0.001.

For urban-captured males, the minimum frequency of song types shared with recordings from the urban field site (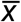= 3.434 log_10_Hz [2761.2 Hz], *N* = 10 song types) did not differ from that of song types not found in field recordings (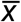 = 3.437 log_10_Hz [2761.6 Hz], *N* = 18 song types; *t* = −1.11, *P* = 0.27). In fact, the means for the minimum frequency of song types were almost identical between those shared and not shared with the urban field recordings (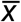 difference = 0.0029 log_10_Hz [0.38 Hz]).

Compared to field recordings from their respective populations, common garden males sang with significantly lower minimum frequencies (mountain, *t*_16.7_ = −4.33, *P* < 0.001; urban, *t*_36.4_ = −8.08, *P* < 0.001; Fig. 3A). In contrast, mountain-captured males sang at significantly higher maximum frequencies in the common garden when compared to field recordings of mountain males (*t*_17.9_ = 2.44, *P* = 0.03; Fig. 3C), but they did not differ in peak frequency (*t*_18.4_ = 1.66, *P* = 0.11; Fig. 3B). The maximum frequency of urban-captured males did not differ statistically from field recordings of urban males (*t*_31_ = 1.65, *P* = 0.10; Fig. 3C), but they did sing at significantly higher peak frequencies (*t*_32.4_ = 2.48, *P* = 0.02; Fig. 3B).

**Figure 3.**
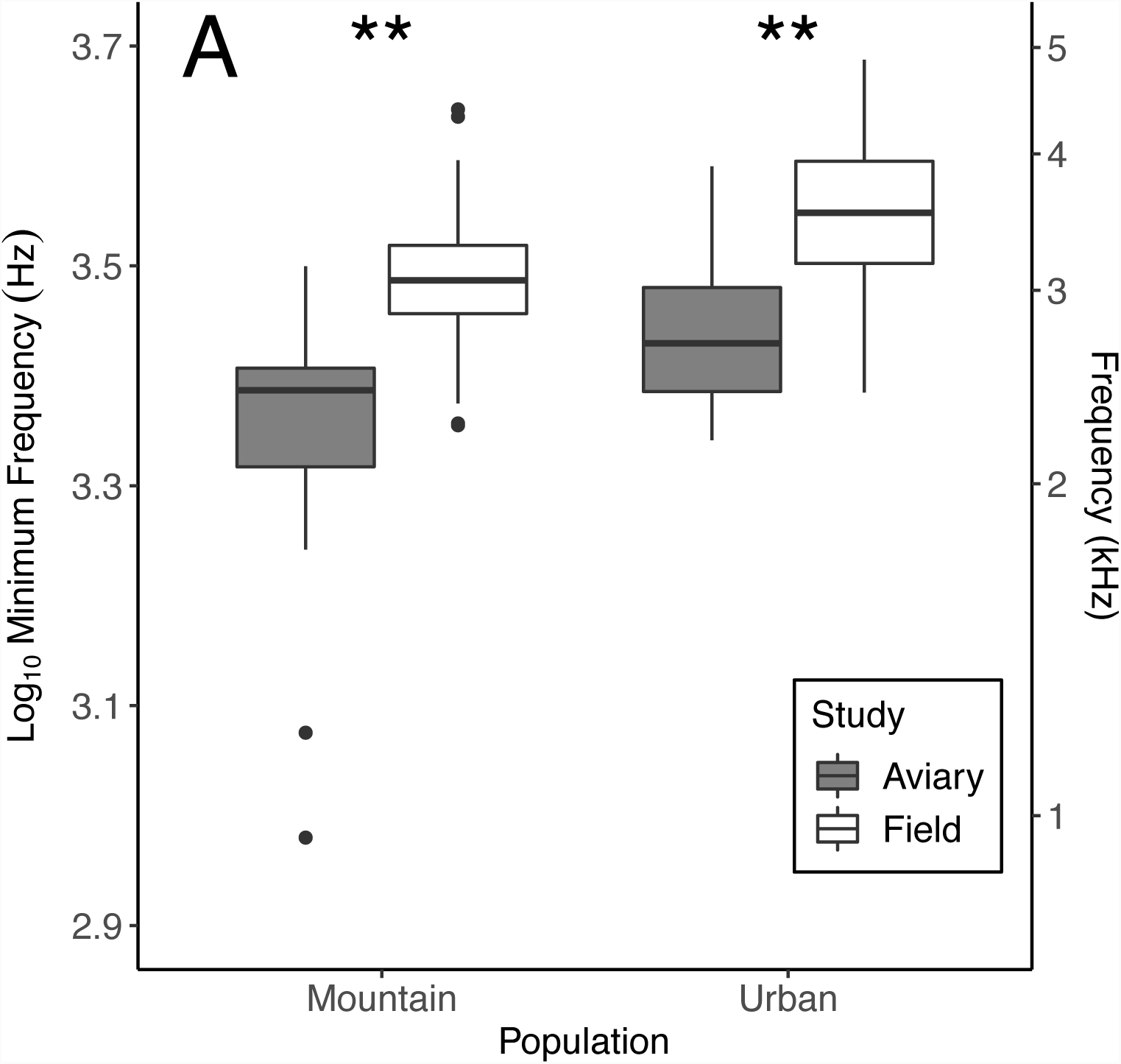
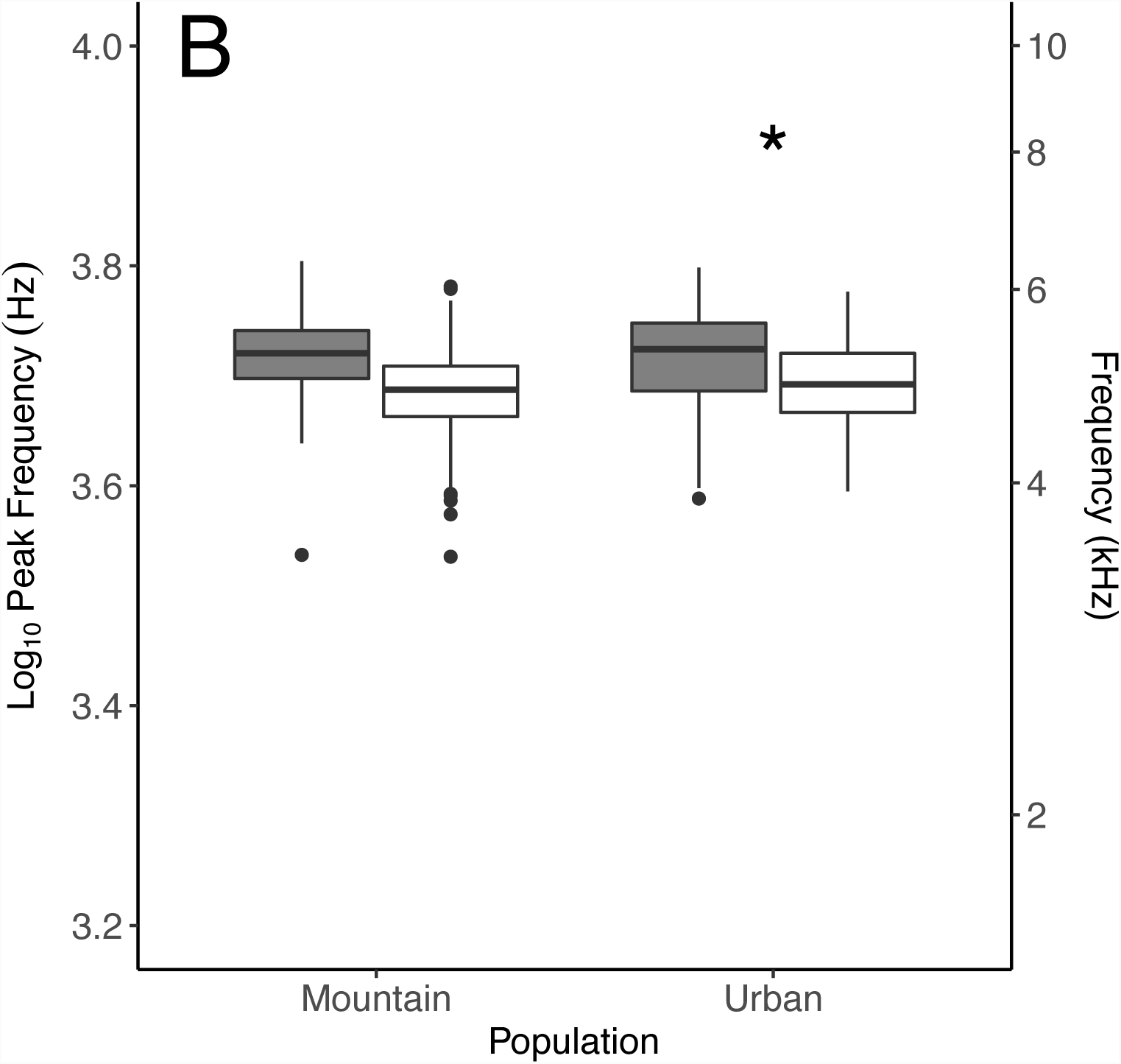
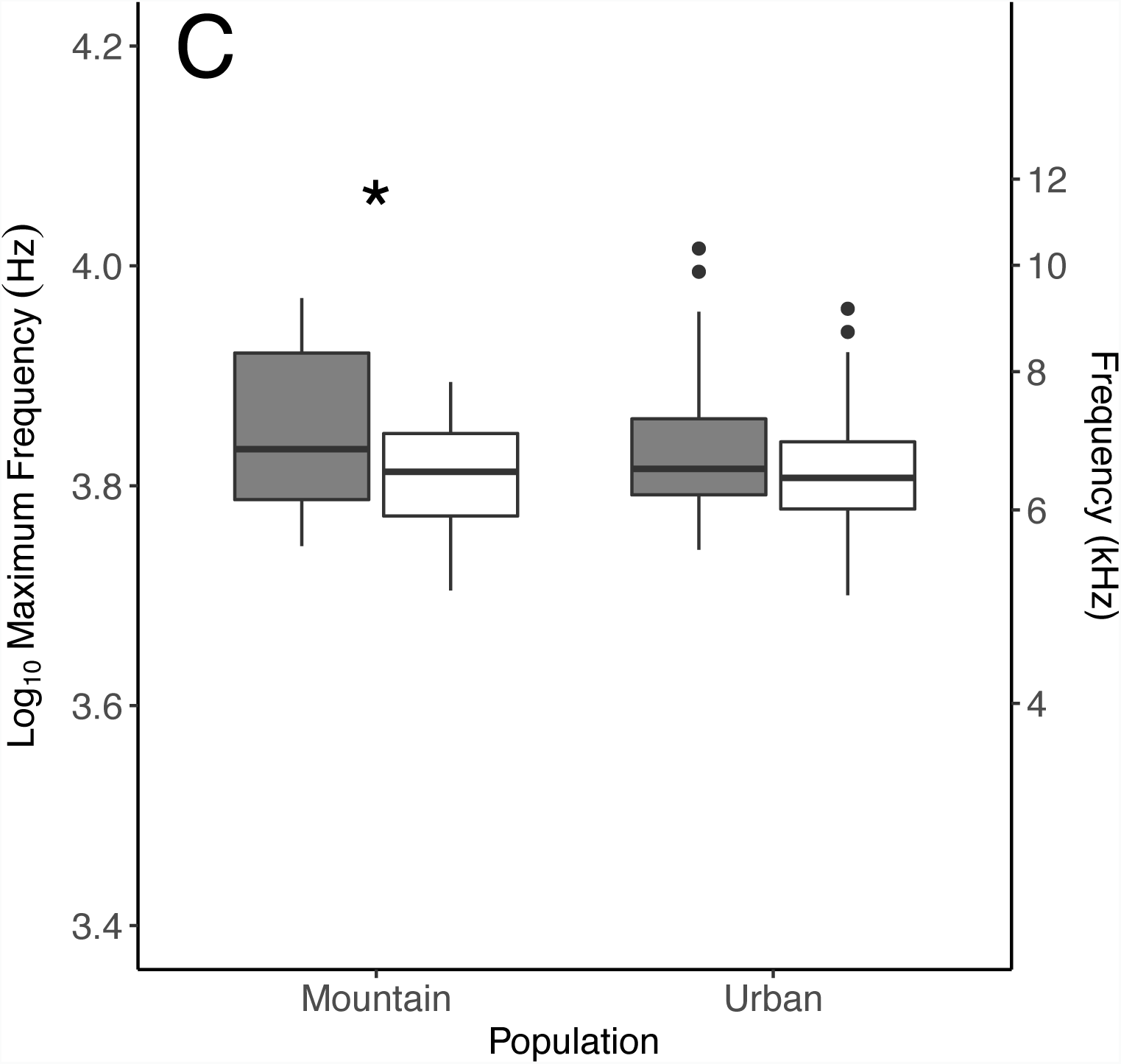
Comparison of the minimum (A), peak (B), and maximum (C) frequency of songs produced by mountain and urban males in the field [19] and in a common garden environment. Boxplots as in Fig. 2. Legend is identical for A-C. \*\**P* < 0.001. \**P* < 0.05.

## Discussion

Mirroring differences observed in the field [19], urban male juncos captured early in life sing at significantly higher minimum frequencies than mountain-captured juncos when both are held in a quiet, common garden environment. This result supports the prediction of the early experience and evolutionary change hypotheses and indicates that the higher minimum frequency of the urban junco population is established early in life, through genetic and/or cultural mechanisms, rather than occurring as a result of behavioral plasticity in response to noise. No significant differences were found in maximum or peak frequency between the two populations in the common garden, indicating that acoustic adaptation in the urban population has acted predominantly on minimum frequency rather than on the frequency of the entire song. We also identified multiple shared songs between the common garden and natal field populations, indicating that early song learning from tutors in the field before capture could contribute to the differences observed in the common garden.

The songs of oscine birds develop through a combination of cultural transmission and genetically-based influences on the morphology and physiology of the vocal production apparatus, on the auditory system, and on learning preferences [63, 64]. While genetic evolution relies on selection or drift acting on standing genetic variation and random mutations, cultural evolution allows for selectively learning pre-existing memes (i.e. cultural selection) or generating novel memes [i.e. cultural mutation] in a non-random, adaptive way [19]. For example, white-crowned sparrows in noisy environments were shown to preferentially learn higher frequency songs and also to elevate the frequency of the learned songs above those of their tutor [21]. A similar combination of cultural selection and cultural mutation has been inferred for urban dark-eyed juncos based on population comparisons of song type meme pools [19].

As in other songbirds, dark-eyed juncos partially rely on conspecific tutors early in life to develop species-typical songs [65], but the exact duration of this sensitive period for song learning is not known. We observed some song type sharing between the common garden birds and field recordings from their natal populations, suggesting that song learning occurred in the field before capture at around 25 to 40 days of age. This timeframe is consistent with the timing of the sensitive period in other closely-related sparrows [66, 67]. However, the majority of song types in both common garden populations were not shared with any known field tutors despite our extensive catalog of field recordings, particularly in the spatially-confined, urban population [19]. In addition to cultural transmission from adult tutors, dark-eyed juncos are known to experience frequent cultural mutations in the form of modifications to learned song types [i.e. improvisation, 68] or *de novo* creation of new song types (i.e. invention, [68]; reviewed in [50]). These frequent cultural mutations likely explain the low incidence of song sharing between the common garden and field populations.

We also observed frequent song sharing among common garden males (41% of song types for the mountain-captured birds, and 29% for the urban-captured) at a much higher rate than typically found in the field, where most neighboring males do not share song types [50–52]. This disparity suggests that much of the song development in the common garden was strongly influenced by peers rather than by adult tutors in the field before capture. The importance of peer interactions is consistent with a previous experiment that showed that when young dark-eyed juncos are reared together without adult tutors they are stimulated to create novel sounds (cultural innovation), copy them from each other, and modify them (cultural improvisation) into a species-typical song [65]. This type of cultural mutation would likely be biased towards higher frequencies if it occurred in a noisy urban environment [19, 21], but our common garden environment was quiet, and, therefore, the direction of this type of cultural mutation should be random or even biased towards low frequencies. Accordingly, we found that juncos from both populations in the common garden sang at significantly lower minimum frequencies than field recordings from their natal populations. This difference between the field and common garden juncos suggests some plasticity in song development, likely related to the quieter acoustic environment in the common garden.

Importantly, all of the changes in song from the wild to the common garden (lower minimum frequencies, learning from peers) did not erode the population difference in minimum frequency. The difference in minimum song frequency between mountain- and urban-captured birds in the common garden was large (417 Hz), and close to the difference reported between the wild populations (540 Hz; [19]). This outcome suggests that the two populations now differ genetically in traits that influence minimum song frequency, thus maintaining the population difference even in the face of an overall decrease in minimum frequency by both common garden populations. The divergent traits that may be responsible for the persistent population difference could be cognitive, such as learning or singing preferences (e.g. the genetic song template [63, 69–71]), or even anatomical or physiological traits that affect song production (e.g. body size [39, 72], or bill morphology [41]). Morphologically-mediated population differences in sound frequency are perhaps less likely because, although urban juncos are slightly smaller than mountain juncos, there is no detectable relationship between body size and song frequency in either of our field populations [73]. Whether the genetic song template or other aspects of neuroanatomy have diverged between the urban and mountain juncos is unknown and represents an intriguing direction for future research.

The mechanisms underlying the evolutionary change and early experience hypotheses are not mutually exclusive and may even reinforce each other. For example, besides the difference in minimum song frequency, the urban junco population studied here also diverged in morphological, reproductive and endocrinological traits [44-47, 74]. Some of these traits appear to have changed by a combination of phenotypic plasticity, which provides an immediate and approximate adaptation to the urban environment, and then selection causing genetic assimilation and the adjustment of the plastic response [47, 75]. Song traits, including minimum frequency, are also likely to undergo such synergy of plasticity and selection. Initially, behavioral flexibility can change songs to provide an immediate reduction of masking by noise [17, 22, 28], and this plasticity simultaneously creates cultural models for which genetically-based learning preferences may be selected upon. Interestingly, song types of urban-captured males that were shared with field recordings, and thus likely to have been memorized from tutors in the urban environment, had an identical average minimum frequency to the unshared song types, many of which would have developed later in the common garden. This result suggests that cultural learning early in life is not the most important explanation for the persistent population difference in song frequency. Instead, around 30 years after colonization of the urban environment [44, 45], it seems likely that the population difference in song frequency is already substantially genetically ingrained.

Broadly, our results suggest that urban environments, and particularly urban noise, may drive the evolution of higher minimum frequencies through a combination of cultural and genetic changes. The urban junco population studied here experienced one of the largest documented increases in minimum song frequency in less than 30 years, indicating that if evolutionary changes are the primary driver, they can happen relatively rapidly [19, 48]. Although behavioral flexibility may provide an immediate escape from masking by environmental noise [17, 22, 28], a combination of cultural evolution and genetic selection on song-related traits can potentially drive more permanent shifts in minimum frequency in chronically noisy environments [20]. The juncos in this study experienced less than 40 days of life with adult song tutors and the noise present in their natal environments, which also suggests that possible developmental mechanisms were triggered in very early life (e.g. memorization of song types, experiencing noise) and had lasting effects. It remains unclear whether the persistent frequency differences between the urban and mountain juncos are driven by experience related changes in song, such as cultural transmission or experiencing noise early in life, or by genetic divergence in traits that influence song learning or production. Future work can begin to disentangle these effects by cross-fostering or hand rearing young birds from urban and wildland environments and tutoring with a wide range of song frequencies.

## Acknowledgments

We thank R. Kiley for assistance with collecting song recordings and H. Hoekstra, R. Lande, K. Marchetti, and the Descanso Ranger District of the Cleveland National Forest for logistical support at field sites.

---

**Table S1.** Results of linear mixed model analyses comparing frequency characteristics between common garden populations measured from power spectra, rather than from spectrograms, using song type as a random effect.

| Song Measurement | Estimate | *t* | *P*-value | N |
| --- | --- | --- | --- | --- |
| Minimum Frequency | 0.081 | 4.528 | **<0.001** | 64 |
| Maximum Frequency | -0.001 | -0.083 | 0.934 | 55 |

**Table S2.** Results of linear mixed model analyses comparing frequency characteristics between common garden populations measured from spectrograms using individual as a random effect rather than song type.

| Song Measurement | Estimate | *t* | *P*-value | N |
| --- | --- | --- | --- | --- |
| Minimum Frequency | 0.095 | 3.011 | **0.003** | 65 |
| Maximum Frequency | -0.025 | -1.437 | 0.151 | 65 |
| Peak Frequency | 0.012 | 0.429 | 0.668 | 65 |

**Table S3.** Results of linear mixed model analyses comparing frequency characteristics between common garden populations measured from power spectra, rather than from spectrograms, using individual as a random effect rather than song type.

| Song Measurement | Estimate | *t* | *P*-value | N |
| --- | --- | --- | --- | --- |
| Minimum Frequency | 0.078 | 2.123 | **0.034** | 64 |
| Maximum Frequency | -0.013 | -0.722 | 0.471 | 55 |

**Figure S1.**
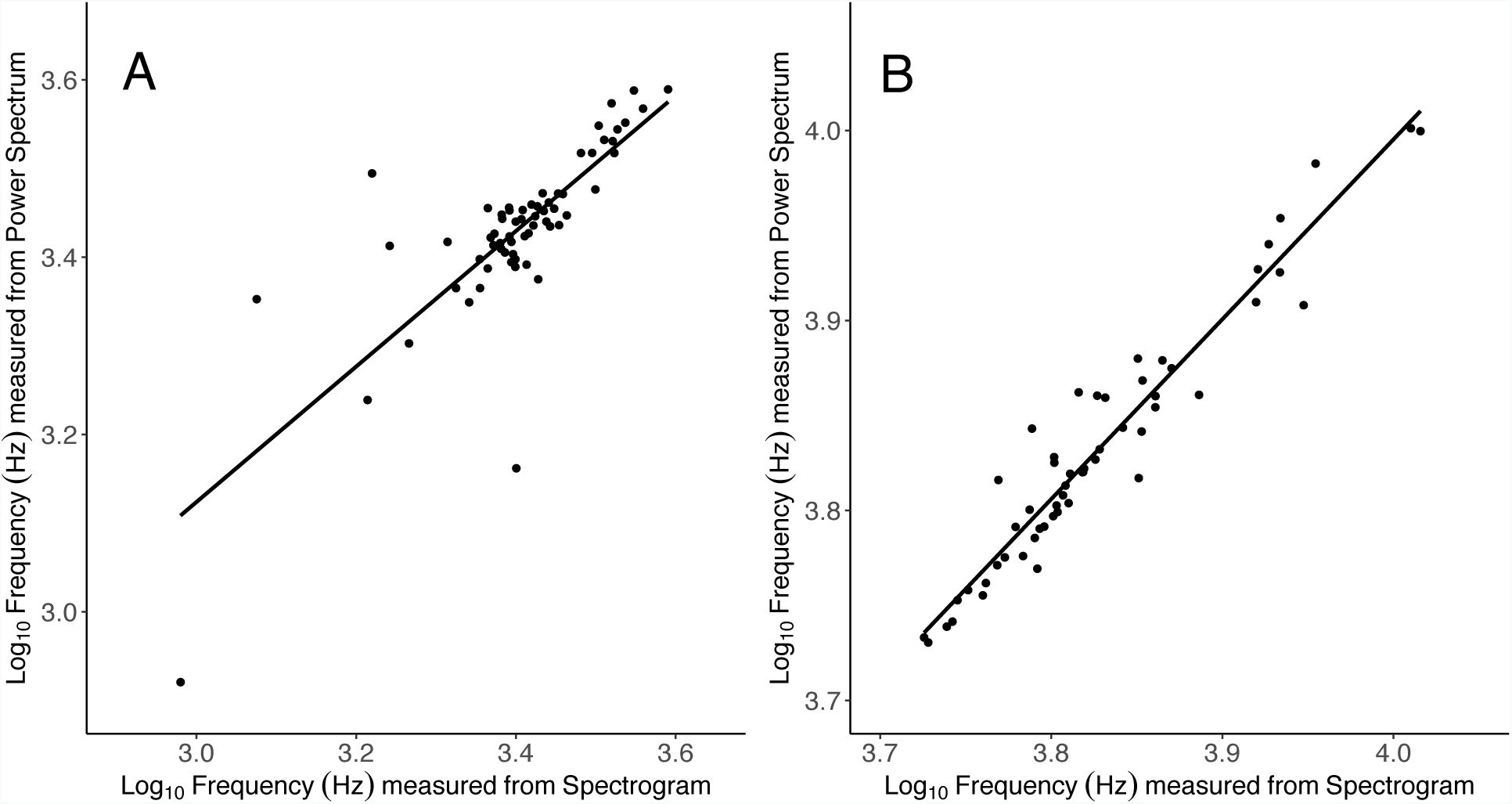
Correlation of (A) minimum and (B) maximum frequencies measured visually from the spectrogram and using a minus 30 dB threshold from the power spectrum.

**Figure S2.**
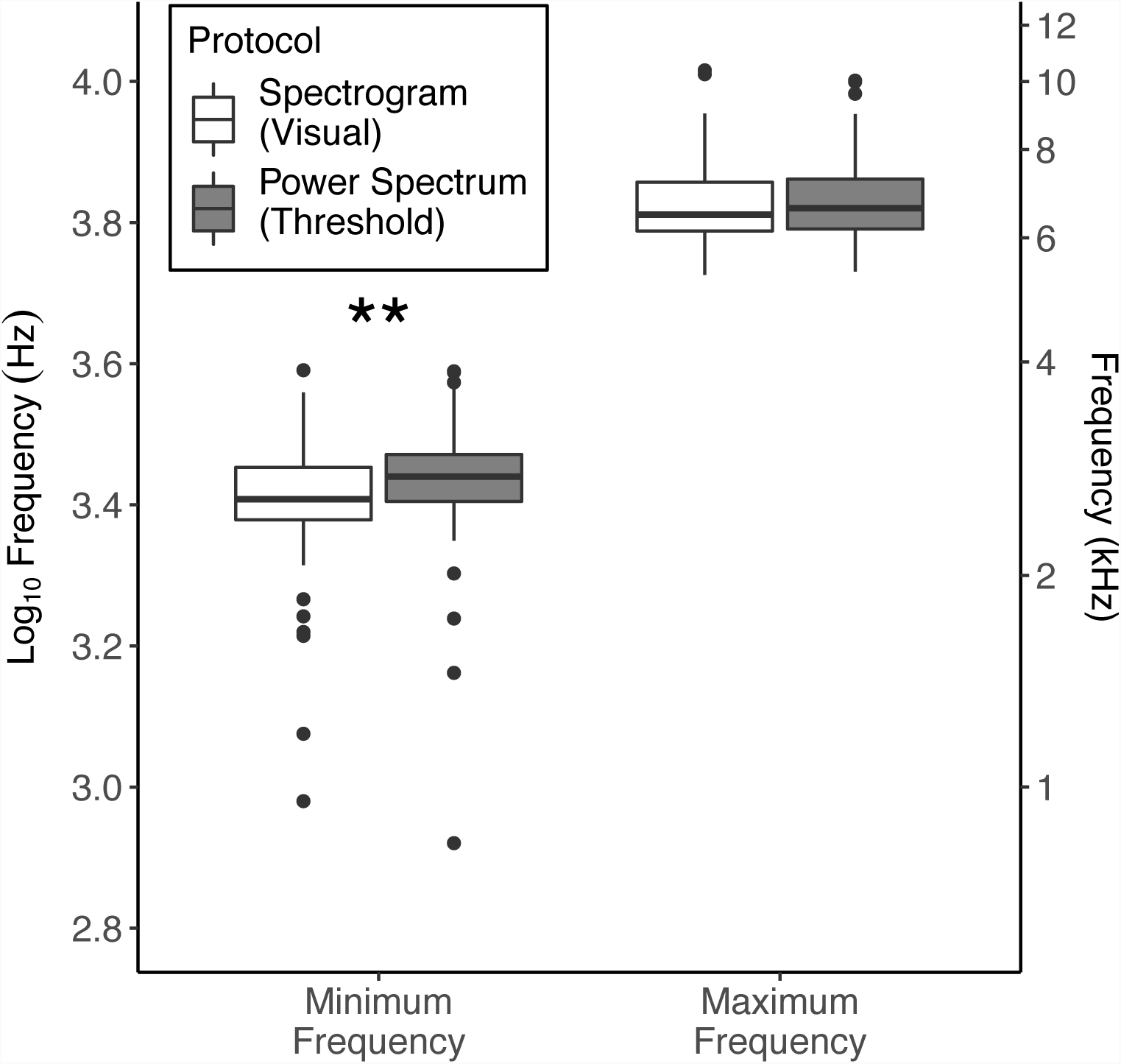
Comparison of the spectrogram and power spectrum techniques for measuring minimum and maximum frequency of songs recorded in the common garden environment. Each box represents the interquartile range and median, whiskers represent range of data within 1.5 times the interquartile range, and dots represent data points exceeding that range. \*\**P* < 0.001

